# TahcoRoll: An Efficient Approach for Signature Profiling in Genomic Data through Variable-Length *k*-mers

**DOI:** 10.1101/229708

**Authors:** Chelsea J.-T. Ju, Jyun-Yu Jiang, Ruirui Li, Zeyu Li, Wei Wang

## Abstract

*k*-mer profiling has been one of the trending approaches to analyze read data generated by high-throughput sequencing technologies. The tasks of *k*-mer profiling include, but are not limited to, counting the frequencies and determining the occurrences of short sequences in a dataset. The notion of *k*-mer has been extensively used to build de Bruijn graphs in genome or transcriptome assembly, which requires examining all possible *k*-mers presented in the dataset. Recently, an alternative way of profiling has been proposed, which constructs a set of representative *k*-mers as genomic markers and profiles their occurrences in the sequencing data. This technique has been applied in both transcript quantification through RNA-Seq and taxonomic classification of metagenomic reads. Most of these applications use a set of fixed-size *k*-mers since the majority of existing *k*-mer counters are inadequate to process genomic sequences with variable-length *k*-mers. However, choosing the appropriate *k* is challenging, as it varies for different applications. As a pioneer work to profile a set of variable-length *k*-mers, we propose TahcoRoll in order to enhance the Aho-Corasick algorithm. More specifically, we use one bit to represent each nucleotide, and integrate the rolling hash technique to construct an efficient in-memory data structure for this task. Using both synthetic and real datasets, results show that TahcoRoll outperforms existing approaches in either or both time and memory efficiency without using any disk space. In addition, compared to the most efficient state-of-the-art *k*-mer counters, such as KMC and MSBWT, TahcoRoll is the only approach that can process long read data from both PacBio and Oxford Nanopore on a commodity desktop computer. The source code of TahcoRoll is implemented in C++14, and available at https://github.com/chelseaju/TahcoRoll.git.

## 1 Introduction

High-throughput sequencing (HTS) has been the leading technology to study genome-wide analyses. A variety of applications have been widely used, including DNA sequencing (DNA-Seq) for characterizing human genetic variation, RNA sequencing (RNA-Seq) for studying gene expression patterns, bisulfite sequencing (Bisulfite-Seq) for discovering epigenetic marks of DNA methylation, and metagenomic sequencing for quantifying the microbial biodiversity in environmental samples or animal body sites. This technology reads out the sequence of DNA fragments and generates millions of reads. The knowledge extracted from these reads allows researchers to answer certain biological questions of interest. To decipher the relevant knowledge from sequencing data, a series of lightweight algorithms has recently been proposed. These algorithms use the notion of *k*-mers for data analysis, which have been claimed to be more efficient and accurate.

*k*-mers are short consecutive subsequences of a genomic sequence. They represent certain signatures to describe different genomes or different regions in one genome. Instead of alignment, existing lightweight approaches pre-compute a searchable database of *k*-mers representing the “signatures,” and count the occurrences of these signatures in sequencing data. RNA-Seq and Metagenomic sequencing are the predominant fields that use *k*-mer approaches. To name a few methods, Sailfish [28], RNA-Skim [35], and Kallisto [5] are the prevalent methods for RNA-Seq transcript quantification; LMAT [2], Kraken [33], and CLARK [26] present efficient strategies to assign taxonomic labels for each metagenomic read. Other *k*-mer applications include studying the CpG island evolution in mammalian genomes using *k*-mer and *k*-flank patterns [6], comparing *k*-mer profiles of family trios to detect disease-causing variants [31], and mining differentially occurred *k*-mers between cases and controls for association mapping [29]. Most of these applications employ a set of fixed-size *k*-mers; however, selecting the appropriate *k* is challenging. The best *k* to characterize different genomic regions can vary. Chae *et al.* [6] have shown that it is necessary to consider patterns of 3- to 10-mers to construct the phylogenetic tree. Rahman *et al.* [29] have proposed to merge the differential occurred *k*-mers to form longer and variable-length sequences for downstream analysis. Ju et *al.* [16] have also demonstrated the advantage of using variable-length *k*-mers for transcript abundance quantification. Yet, existing *k*-mer counters are optimized to process *k*-mers of a fixed length. Inevitably, it is critical to have a feasible data structure to store *k*-mers of different sizes, and to analyze sequencing data efficiently and accurately.

Given a set of *k*-mers with the same size, a straightforward counting method is to index them with a hash table and scan through read sequences with a fixed-size window. If a set contains *k*-mers of different sizes, the read sequences need to be scanned multiple times with different *k*’s. This repetition limits the analysis to evaluate only a small range of *k*’s. An alternative approach is to use existing efficient *k*-mer counting algorithms. Read sequences are first indexed and stored as *k*-mers, and the number of occurrences of each *k*-mer can be computed from the index. One of these counters, **Jellyfish** [22], has been widely used as the underlying structure for Sailfish, Kraken, DIAMUND [31], and hAwK [29]. **Squeakr** [27] is a newly developed algorithm that also employs the thread-safe approach to efficiently query the counts of a specific *k*-mer. In addition, probabilistic data structures are also commonly used in *k*-mer counting. Its implementations include **BFCounter** [24] and **khmer** [34]. Disk-based hashing is another popular hashing technique, and its related algorithms are **DSK** [30], **KAnalyze** [4], **MSPKmerCounter** [20], and **KMC** [11, 12,18]. Other data structures to facilitate *k*-mer counting are burst tries used in **KCMBT** [21], and suffix-array like structures employed in **Tallymer** [19] and **MSBWT** [15]. Several of these implementations, such as BFCounter, khmer, and MSPKmerCounter, restrict the choice of *k* to fall within a certain threshold to mitigate the memory consumption and running time. Suffix-array based approach is the only one that presents the potential to process *k*-mers of variable lengths. All other methods are designed to process sequences with a fixed *k*. Thus, repeating the indexing step for different *k*’s is unavoidable. It is worth mentioning that SBT [32] is a hybrid approach that uses Jellyfish as the underlying structure, bloom filter to record frequently appeared *k*-mers, and a tree structure to store a collection of bloom filters. It is designed to query frequently appeared sequences in multiple experiments, such as expressed genes in RNA-Seq. Its aim is different than the problem we formulated as their signature sequences can be longer than reads, and report approximate count for each *k*-mer used in indexing. Therefore, it is not included in our comparison. A detailed summary of the related approaches is described in Section C in the supplementary materials.

Computing the frequency of a set of *k*-mers with different sizes can be reduced to a multiple pattern matching problem [25] in computer science. A linear solution to scan the reads once is the Aho-Corasick algorithm [1], which constructs a tree automaton upon the trie of keywords. In this tree, there are additional links between internal nodes to facilitate the *k*-mer matches without the need for backtracking, i.e., jumping back and forth of the query sequence. A drawback of having this automaton is the memory requirement for storing long or large numbers of *k*-mers. As we increase the number or the length of *k*-mers, the tree grows wider and deeper respectively. In addition, larger *k*-mers are usually more diverse and have shorter common prefixes, requiring more space for *k*-mer representation. Thankfully, the concise representation of DNA molecules allows further reduction in memory requirement of this automaton. Since these *k*-mers are composed of only four different characters: `A`, `C`, `G`, and `T`, they can be succinctly represented in a binary format. We propose to use one bit, i.e., 0 or 1, to represent these four characters while constructing the automaton. This binarized representation allows us to significantly shrink the structure of the tree, and to substantially reduce the memory usage. To avoid collisions of *k*-mers with identical binarized representations on the tree, each node contains a hash table to recover the original *k*-mers.

It is important to note that the goal of our work is not to compute the frequency of all possible *k*-mers with different sizes, but to profile a pre-defined set of variable-length *k*-mers as signatures in sequencing reads. An example of a pre-defined set can contain the genetic markers of different microorganism in metagenomic studies. Since the term signatures here refer to a set of representative *k*-mers, we use these two terms interchangeably throughout this paper. Our contributions are fourfold. First, we emphasize the needs of having a viable data structure to store and to profile *k*-mers of different sizes in DNA sequences. Second, to the best of our knowledge, this is a pioneer work to profile a vast amount of variable-length *k*-mers simultaneously in genomic data. We propose to apply the Aho-Corasick algorithm with a memory efficient automaton. Third, we leverage the properties of DNA sequence to construct an efficient in-memory structure, and employ the rolling hash technique to accelerate the match. Fourth, we adapt existing *k*-mer counters to perform the same task and conduct a comprehensive analysis of 14 different methods. Results show that our method, TahcoRoll, is more efficient in profiling signatures with a wide range of sizes than conventional *k*-mer counters. It is also resistant to the change of read lengths and number of reads. Most importantly, TahcoRoll is able to analyze long read data from both PacBio and Oxford Nanopore in a desktop computer while other approaches fail.

## 2 Methods

In this section, we propose the <u>t</u>hinned <u>Ah</u>o-<u>Co</u>rasick automaton accelerated by <u>roll</u>ing hash (TahcoRoll) to profile variable-length *k*-mers in genomic data.

### 2.1 Problem Statement

We first formally define the problem of signature profiling. In this paper, we focus on counting the occurrences of a set of representative *k*-mers instead of all possible *k*-mers because of the following two reasons. First, the number of all variable-length *k*-mers in a DNA sequence could be too enormous to be verified. More specifically, the lower-bound of time complexity to examine all occurrences of possible *k*-mers with different *k* is at least *O(L^2^)* for a read of length *L*. Second, there is no need to check all possibilities when a signature set is given.

Suppose that ***P*** is the set of representative *k*-mers as signatures, where the length *k* could be different across the set. Given a set of sequencing reads ***T***, our goal is to profile the occurrences of each signature *p* ∊ ***P***. Note that the lengths of occurred patterns are shorter than the read length.

More formally, for each signature *p* ∊ ***P***, we aim to develop an efficient algorithm to compute the number of overall occurrences *c_p_* as follows:

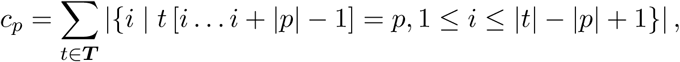

where |*p*| and |*t*| indicate the lengths of the signature *p* and the sequencing read *t*, and *t* [*i* … *j*] denotes the substring of *t* from the *i*-th to the *j*-th character. Figure 1 shows an example with five signatures ***P*** = {`ATT`, `GA`, `TTG`, `AGAT`, `TC`} and two sequencing reads **T** = {AATTGAGAT, ATTGACATCG}. Each segment indicates an occurrence in the read for the corresponding signature. Occurrences could overlap with each other. A signature could have multiple occurrences in a single read (e.g., `GA`) or across a set of reads (e.g., `ATT`). For each signature *p* ∊ ***P***, occurrences in the read set ***T*** are counted as a number *c_p_,* which is the objective of signature profiling.

**Fig. 1:**
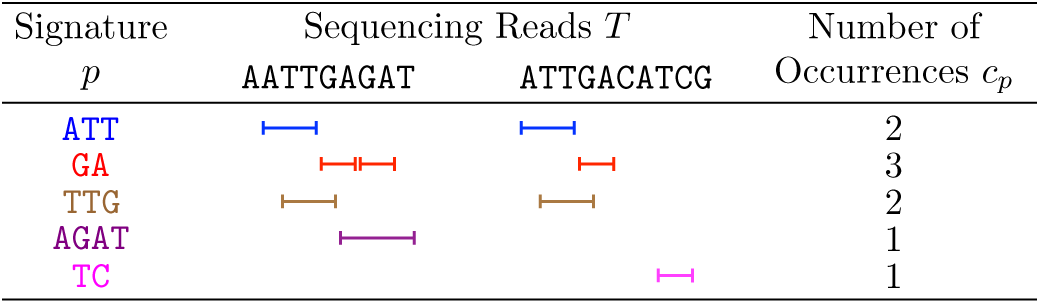
An example with five signatures and two sequencing reads in signature pro_ling. Each segment represents an occurrence of the corresponding signature in the read.

#### 2.2 Aho-Corasick Automaton

To solve signature profiling, we reduce the task to multiple pattern matching [25] by mapping signatures onto patterns and reads onto the input text to be matched. The algorithms of multiple pattern matching find all occurrences in a read for each signature, so the results of signature profiling can be obtained by aggregating these occurrences. As the pioneer study, we propose to apply the Aho-Corasick algorithm (AC) [1], which is one of the state-of-the-art approaches for multiple pattern matching, to profile signatures.

AC conducts the matching process along a trie that corresponds to patterns. Each node in AC has a failure link that allows fast transitions from one node to the other representing its longest possible suffix without backtracking. Informally, AC constructs a finite state machine (or an automaton) that resembles a trie and failure links. The pattern matching process can be treated as transitions between nodes in the automaton, and failure links provide efficient transitions between failed matches. Figure 2 shows an example of AC with five signatures. Black solid links are trie lines, and red dashed lines are failure links. For example, the node of `AGAT` has a failure link to the node of `AT`. Given a sequencing read `ATTTG` to be profiled, AC will first match the signature `ATT` in the *blue* node. Then, it fails to match the third `T` and transits to the *orange* node that still has no child of `T`. After traveling along the failure link again to the *yellow* node, both the last two characters `TG` can proceed towards the *orange* and *brown* nodes that indicate a match of the signature `TTG`.

**Fig. 2:**
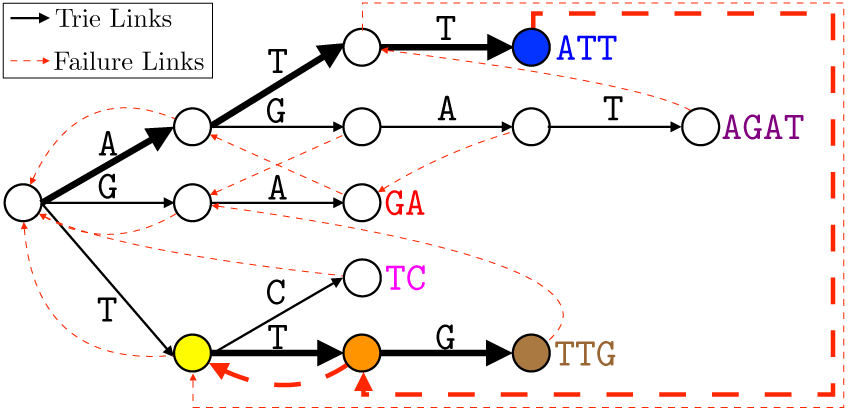
An example of the AC automaton with five signatures. Black solid lines are trie links, and red dashed lines are failure links. Colored nodes and thicker lines are important nodes and paths while profiling the read ATTTG.

The construction of the automaton in AC with signatures *p* ∊ ***P*** only requires a simple breadth- first search (BFS) with a 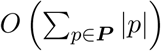 linear time complexity. To profile signatures in reads *t* ∊ ***T***, AC only needs to simulate transitions on the automaton, which also has a linear time complexity 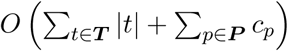. The space complexity of AC is also linear, 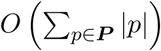, to maintain a node and a constant number of links for each character in signatures. Therefore, in theory, AC is a perfect fit to deal with signature profiling.

#### 2.3 Thinned Automaton with Binarized Pattern Matching

Even though we have shown the theoretical capability of AC for signature profiling, there are still some hurdles in practice. One of the most significant problems is the memory usage when the number of signatures is enormous. More specifically, each individual character in signatures can be referred to as a trie node, which provides plenty information and consumes a considerable amount of memory. For example, the plain AC requires more than 240 GB of memory to process 24 millions of signatures whose lengths range from 135 to 151. Especially for the cases of signatures with fewer and shorter common prefixes, nodes tend to have more child nodes, and the greater width leads to the increase of memory usage.

To reduce both the number of nodes and the width of the automaton, we propose the thinned automaton with binarized pattern matching. More formally, each signature *p* [1… |*p*|] ∊ ***P*** is transformed into a binarized pattern *p′* [1… |*p*|] before being added into the automaton. The **i**-th character *p′* [*i*] of *p′* is defined as follows:

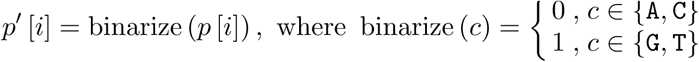

Note that these four characters can be randomly divided into two groups. Compressing two characters into one bit 0 or 1, binarized patterns improve the representation capability of a depth-*d* node in the trie from 1 to 2^*d*^ unbinarized pattern(s), thereby reducing both the width of the automaton and the number of nodes. In this paper, the automaton with binarized patterns is named *thinned automaton* because of its reduced width. Here, we conduct a theoretical analysis of the improvement of the thinned automaton against the plain AC. For convenience, we assume that each character in a signature is uniformly distributed. To estimate the worst-case scenario, we assume that every signature has the largest length *m* observed in the set. While inserting a signature into a trie, the number of newly added nodes depends on the presence of its prefixes in the trie. Proposition 1 gives an expectation of finding prefixes for *n* signatures with *c* possible characters.

##### Proposition 1 (Proved in Section A.1)

*Given n signatures with c possible characters to be added into a trie, the expected number of signatures that fail to find their length-i prefixes along the trie during its insertion is 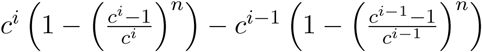, *where* 0 ≤ *i* ≤ *m**.

Based on Proposition 1, the expected number of nodes in a trie can be given by Proposition 2.

##### Proposition 2 (Proved in Section A.2)

*Given n signatures of length m with c possible characters to be added into a trie, the expected number of trie nodes is 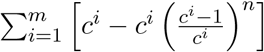.*

Following Proposition 2, Proposition 3 derives the expected improvement in the number of trie nodes when the number of signatures is sufficient

##### Proposition 3 (Proved in Section A.3)

*When the number of signatures in the automaton is approaching a large number, the expected number of nodes in the thinned automaton is only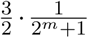of those in the plain AC.*

As shown in Proposition 3, the improvement with the thinned automaton is guaranteed under the assumption mentioned above. However, DNA sequences are known to be biased. In the scenario like this, where the characters of each signature are not uniformly distributed, the improvement can only be more pronounced because more duplicated segments lead to fewer trie nodes. Moreover, the longer the signatures, the more significant the improvements.

Even though the thinned automaton reduces the number of nodes, compressed representations may lead to collisions. Figure 3 shows an example of binarized results for five patterns and two sequencing reads, and Figure 4 further illustrates the corresponding thinned automaton. Two signatures `GA` and `TC` share the same binarized pattern `10` and result in a collision when reaching the *yellow* node in Figure 4. Substrings of a read with identical binarized representations may also lead to false alarms. For instance, `ATCG` in the second read, which is not a signature, has the same binarized representation `0101` to the signature `AGAT`. To maintain the correctness of signature profiling, each match to a binarized pattern needs to be verified with the original signatures. On the other hand, it would be very time-consuming if signatures have serious collisions in certain nodes or extensive lengths. More precisely, a naïve comparison costs 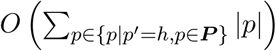 time to verify signatures with the same representation *h*.

**Fig. 3:**
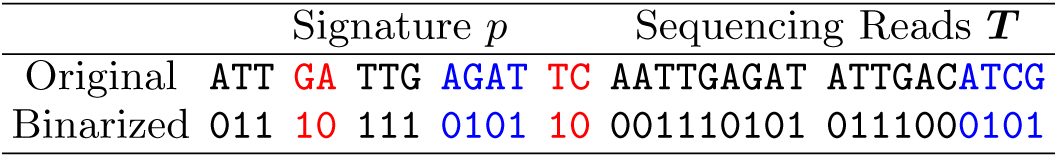
The binarized representations for five patterns and two sequencing reads. Two signatures GA and TC share the same binarized pattern (red). A substring in a sequencing read ATCG has the identical binarized form to the signature AGAT (blue).

**Fig. 4:**
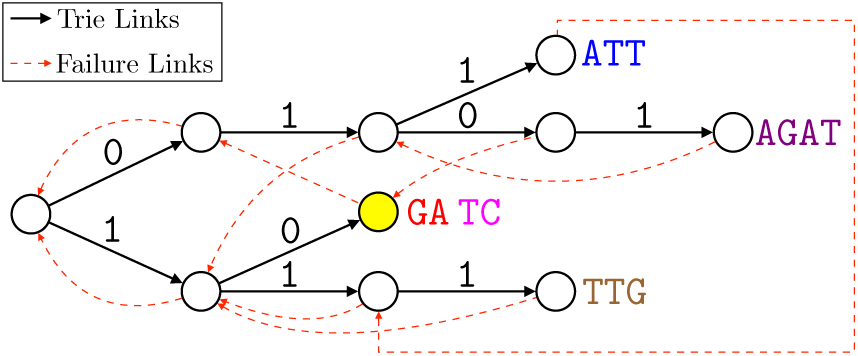
An example of the thinned automaton of AC with five signatures. Black solid links are trie links, and red dashed links are failure links. The *yellow* node represents two signatures GA and TC.

#### 2.4 Acceleration by Rolling Hash

Using hash functions is an intuitive idea to speed up comparisons between strings. As the lengths of signatures can vary, arbitrary length substrings of the read *t* ∊ ***T*** may be required to compute hash values during verification. However, on-the-fly computation of hash values takes an additional linear time *O* (|*t*|) for each checkup; precomputing all possible substrings is also infeasible due to dispensable computations and extensive *O*(|*t*|^2^) additional memory usage.

To accelerate verification, we propose to apply rolling hash [8] that alleviates the time complexity for each checkup from linear to constant with a linear-time preprocessing and an additional linear memory consumption. Rolling hash is a family of hash functions where the input is hashed with a window that moves through the input. A new hash value can be rapidly calculated from the given old hash value in *O*(1) time. It also allows for *O*(1) query time on the hash value of any substring in the input with content-based slicing. In this paper, we apply the Rabin-Karp algorithm [17] to implement a rolling hash function as shown in Section B in the supplementary materials. As a theoretical analysis, Proposition 4 gives a theoretical upper-bound of the collision probability.

#### Proposition 4 (Gonnet and Baeza-Yates [14])

*The probability of two different random strings of the same length having the same hash value in Rabin-Karp rolling hash is P(collision) ≤ 1/q, where q is the prime modulus in computations of the Rabin-Karp algorithm.*

When the prime modulus *q* is large, rolling hash has been shown to provide a small probability of hash collision [14].

To apply rolling hash for acceleration, each node contains a hash table that maps a hash value onto the original signature. When transitioning to a node, the hash value of the matching substring in the read can be rapidly calculated and verified for its presence in the hash table. As a result, the average time complexity of each checkup can be reduced to *O*(1). The overall time complexity of TahcoRoll is 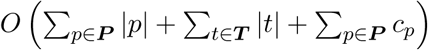, including the construction of the thinned automaton and the matching process. The only memory overhead is hash tables with exactly |***P***| hash values, which is an amortized *O*(|***P***|) space.

## 3 Data

The performance of different algorithms are affected by four factors: signature lengths, number of signatures, read length, and number of reads. To evaluate the running time and memory footprint of different methods, we use synthetic reads and signature sets containing variable-length *k*-mers as described below. The flexibility of synthetic reads allows us to closely examine the effects of these factors. The randomly generated signatures are designed to test the worst scenario as their characters are uniformly distributed and may not share as many common prefixes as in the real sequencing applications. We also generate signatures from both genomic and transcriptomic sequences to analyze real reads from a diverse range of sequencing platforms.

### 3.1 Representative *k*-mers Generation

To examine the effects of signature number and size, we generate four batches of *k*-mers with different ranges of *k*. The first batch, denoted by “small,” contains *k*-mers ranging from 15 to 31 nucleotides. The second batch is considered as the “medium” batch, ranging from 65 to 81 nucleotides. The third batch is designed for longer *k*-mers, denoted by “large”, and their sizes range from 135 to 151. The last batch of *k*-mers ranges from 15 to 151, and is denoted by the “wide” batch. In each batch, there are four sets containing 1.2, 6, 12, and 24 millions of *k*-mers.

The sequence of each *k*-mer is randomly assigned with four characters `A`, `C`,`G`, and `T`, and a random length that falls within the range described above. Each *k*-mer is normalized and represented by its canonical form (i.e., the lexicographical minimum of itself and its reverse complementary sequence). Duplicated *k*-mers are removed from the list.

### 3.2 Synthetic Datasets

We used polyester [13] to generate 13 sets of single-end RNA-Seq reads for selected transcripts. We randomly selected 2–10% of the protein-coding transcripts in each sample. The transcript sequences are based on Ensembl release 81 of Human Genome GRCh38 [10]. Each sample contained 10-115 millions of reads, with read lengths of 75bp, 100bp, 125bp, 150bp, and 180bp.

### 3.3 Real Datasets

We download publicly available datasets generated from a diverse range of sequencing platforms. The first dataset contains two experiments to study the transcriptomic analyses for lymphoblastoid cells [7]: SRR1293901 is a 2×262 cycle run from Illumina MiSeq and SRR1293901 is a 2×76 cycle run from Illumina HiSeq 2000. The second dataset, GSM1254204, aims to characterize the transcriptome of human embryonic stem cells using PacBio long reads [3]. The third datasets are generated by Oxford Nanopore to study the whole genome of the breast cancer model cell line. It contains three datasets, SRR5951587, SRR5951588 and SRR5951600 with different read lengths.

For the RNA-Seq experiments, we use Fleximer [16] to generate a list of 10,962,474 signatures that can characterize different human transcript isoforms. For WGS, a list of 10,935,397 short sequences from the reference genome is randomly selected as the signatures. Since long reads contain a higher error rate, we cannot set the *k* to be too large. Thus, they range from 25 to 60bp.

## 4 Experiments and Results

### 4.1 Software Adaptation

Since most of the *k*-mer counters process reads with a fixed *k*, we adapt each counter to account for a set of variable-length *k*-mers. For each counter, we use a python script as a wrapper to handle different *k*-mer sizes and to call the appropriate functions from the command line. All of the experiments are conducted using a single thread. We include all the related *k*-mer counters mentioned in the Introduction, except Squeakr as it is released less than a month ago, and we do not have sufficient time to properly configure the software for a fair comparison. We implement two baseline methods. The first is a naïve implementation, denoted by “Naïve”. It uses a hash table to store *k*-mers and scans through the reads multiple times with different window sizes. Theoretically, Naïve is light in memory, but requires an extensive running time. The second baseline is a publicly available implementation of the Aho-Corasick algorithm, denoted by “Plain AC”. We describe the configuration details of each method in Section C in the supplementary materials.

### 4.2 Automaton Construction

The memory consumption of AC is sensitive to the composition of signature patterns such as *k*- mer lengths, the number of *k*-mers, and common prefixes shared by different *k*-mers. In our first analysis, we evaluate the effectiveness of the binarized representation of the automaton. Figure 5 compares the time and memory of automaton construction in TahcoRoll against Plain AC over 12 sets of signatures. Our thinned automaton consistently requires less time than the full automaton for construction. As we increase the number of *k*-mers, the construction time rises. The memory of the thinned automaton is significantly reducing to nearly half of the memory required in plain AC. Larger *k*-mers are more diverse in their sequences, and often share shorter common prefixes with others. This phenomenon is reflected in this analysis, where the “large” batches of *k*-mers require more time and memory than others. The “wide” batches of *k*-mers need less computational resources than the “large” ones because their *k*-mer sizes are widely spread from 15 to 151bp.

**Fig. 5:**
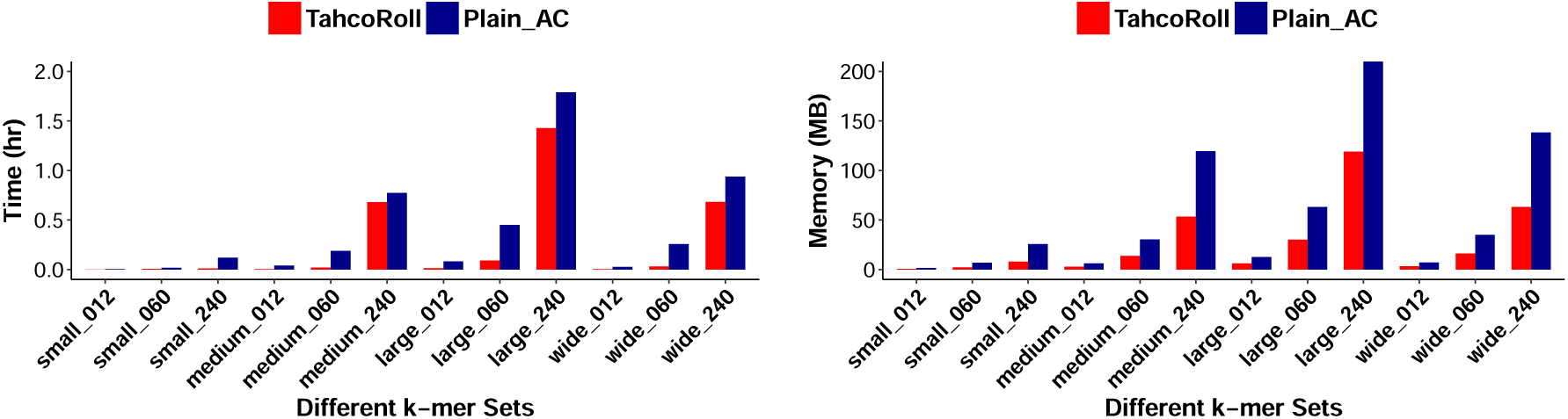
Running time (left) and memory footprint (right) for constructing the keyword automaton given different sets of *k*-mer patterns. A lower value represents a more efficient approach.

### 4.3 Pilot Study of 14 Approaches

We perform a preliminary assessment of the memory footprint and running time on 10 *k*-mer counters, together with two baselines and TahcoRoll. Since most of these counters are designed to process data with a fixed *k*, they present a limitation on handling a broad range of *k*’s. In addition, a few of these algorithms limit the choice of *k*. For these reasons, we inspect the capability of different approaches using small datasets, including three sets of synthetic reads with 75bp and three “small” batches of signatures. Jellyfish provides two ways to retrieve the *k*-mer counts: through its query function (denoted by Jellyfish_query), or through dumping the index to a human readable format (denoted by Jellyfish_dump). We describe them in Section C in the supplementary materials.

We separate the analyses into two panels as demonstrated in Figure 6. The top panel focuses on different numbers of reads, and the bottom panel focuses on different numbers of *k*-mers. Methods in the bottom-left corner of each plot indicate being both time and memory efficient. As we predicted, Naïve uses very little memory, but takes a long time to complete. Plain AC is fast, but requires a large amount of memory as we increase the number of *k*-mers. TahcoRoll is the most efficient approach in five out of these six analyses. KMC performs slightly better than TahcoRoll when there are 24 million *k*-mers, but its memory footprint increases linearly as there are more reads, while the memory usage remains constant in TahcoRoll. Jellyfish_query is memory efficient when processing a small number of reads, but its running time increases significantly with more reads.

**Fig. 6:**
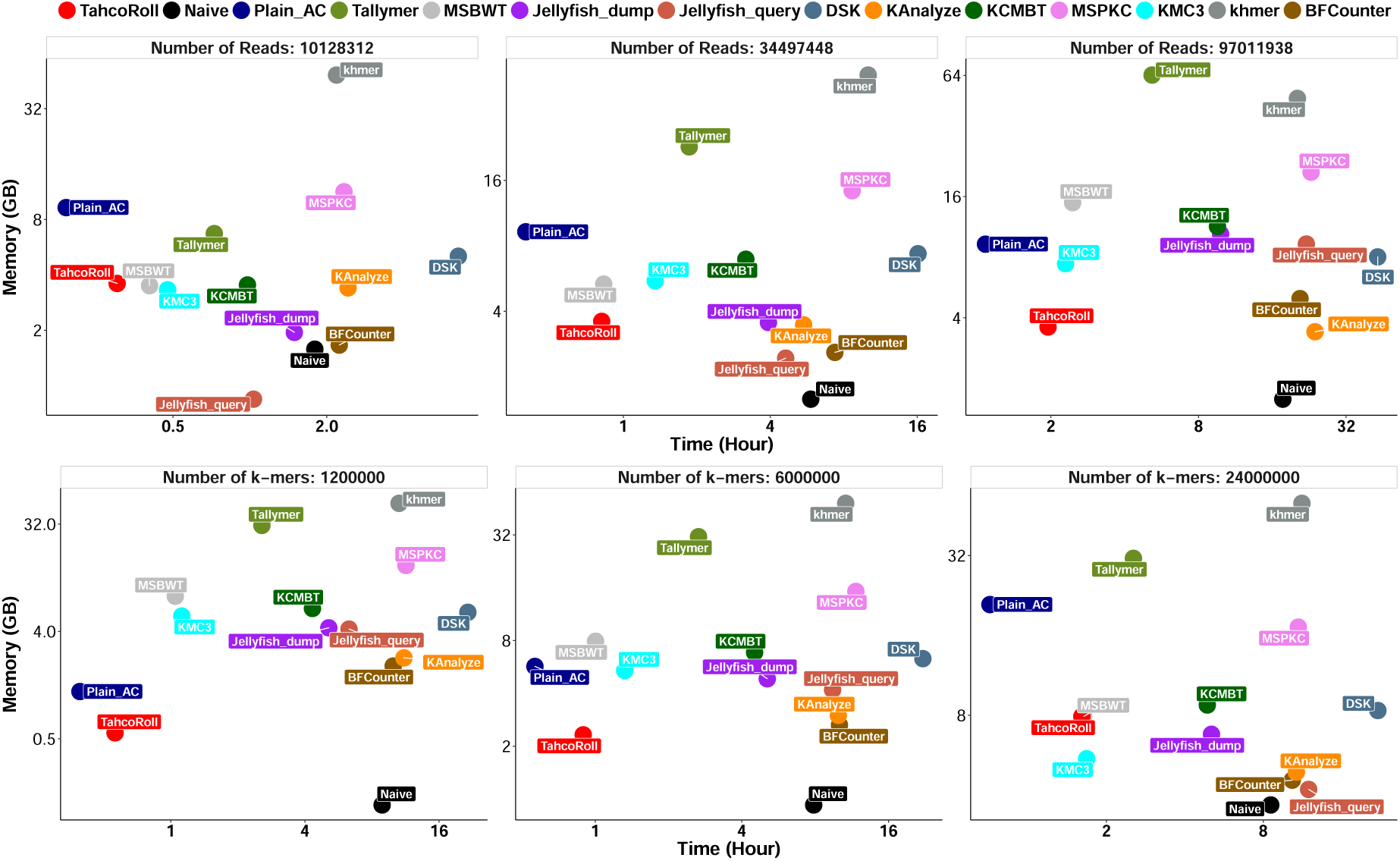
Running time (x-axis) and memory footprint (y-axis) of counting “small” batches of *k*-mers on synthetic reads with 75bp. The top panel examines the effect of the number of reads, and the bottom panel examines the effect of the number of *k*-mers. Each pair of values (running time and memory usage) represent the average of three measurements, either across *k*-mer sets (top) or read sets (bottom). Points located on the lower left corner of each plot indicate more efficient approaches. Jellyfish_query uses less memory when the number of reads is small; KMC3 performs slightly better when there are more *k*-mers. However, both of them do not scale well with more reads.

Using Naïve as a guidance in the running time analysis, DSK, KAnalyze, MSPKC, khmer, and BFCounter are much slower in at least five of these analyses. To avoid waiting on extremely slow counters, we remove these approaches from further analyses. From the memory usage perspective, Tallymer and khmer require much more memory than Plain AC in five of these analyses, which are also eliminated. Although KCMBT is reasonably efficient, it cannot handle *k*-mers larger than 31bp. Thus, it is also excluded from the rest of the experiments. Jellyfish_dump takes slightly more memory than Jellyfish_query, but is much faster with a larger number of reads and *k*-mers. The remaining analyses are carried out for Plain AC, Jellyfish_dump, KMC3, MSBWT, and TahcoRoll.

We also check the accuracy of these counters. We compare the counts reported by each approach to Naïve. As expected, probabilistic approaches, BFCounter and khmer, neglect singleton *k*-mers. All other approaches agree with Naïve. The results are omitted due to space limitations.

### 4.4 Extensive Study on Synthetic Datasets

We use 1.2 million *k*-mers ranging from 15-151bp (wide batch) to evaluate the scalability on read length and number of reads. We highlight the most memory efficient approach of each experiment in Table 1. Jellyfish here refers to jellyfish_dump as described in the pilot study. Since the memory consumption of both TahcoRoll and Plain AC depends on the signature set, they are able to withstand longer reads and a larger number of reads. TahcoRoll uses the least amount of memory across all experiments. We also highlight the top two time-efficient approaches, which are two implementations of the Aho-Corasick algorithms. It is expected that TahcoRoll takes slightly more time than Plain AC as it needs to resolve the collision of *k*-mers with identical binarized representations. We terminate Jellyfish as it runs much longer than three days on datasets with a large number of reads. The memory footprints are recorded at termination.

**Table 1:** Time (Hour) and Memory (GB) of Synthetic Datasets over Different Read Sets

| Read Length | Total Reads | TahcoRoll |  | Plain AC |  | MSBWT |  | KMC |  | Jellyfish |  |
| --- | --- | --- | --- | --- | --- | --- | --- | --- | --- | --- | --- |
|  |  | Time | Mem | Time | Mem | Time | Mem | Time | Mem | Time | Mem |
| 75bp | 10,128,312 | **0.21** | **3.36** | **0.12** | 6.75 | 0.30 | 3.50 | 0.68 | 3.32 | 5.45 | 4.95 |
|  | 34,497,448 | **0.50** | **3.36** | **0.34** | 6.75 | 0.89 | 5.35 | 2.16 | 5.53 | 16.31 | 5.89 |
|  | 97,011,938 | **1.18** | **3.36** | **0.94** | 6.75 | 1.78 | 14.93 | 6.41 | 8.00 | 37.14 | 19.40 |
| 100bp | 11,397,007 | **0.21** | **3.36** | **0.21** | 6.75 | 0.77 | 3.96 | 1.99 | 3.86 | 13.84 | 5.89 |
|  | 41,054,662 | **0.71** | **3.36** | **0.56** | 6.75 | 1.12 | 8.31 | 7.59 | 7.57 | 46.02 | 15.58 |
|  | 114,813,452 | **3.08** | **3.36** | **1.38** | 6.75 | 5.16 | 23.15 | 17.70 | 20.09 | 52.00 | 44.39 |
| 125bp | 10,822,319 | **0.39** | **3.36** | **0.21** | 6.75 | 0.59 | 4.42 | 3.95 | 4.16 | 23.97 | 9.60 |
|  | 107,375,244 | **3.65** | **3.36** | **1.57** | 6.75 | 4.18 | 26.79 | 14.13 | 34.59 | >72.00 | >41.29 |
| 150bp | 27,628,054 | **1.27** | **3.36** | **0.56** | 6.75 | 2.41 | 8.25 | 16.19 | 14.87 | 6.00 | 33.05 |
|  | 57,437,772 | **1.96** | **3.36** | **1.22** | 6.75 | 3.90 | 17.08 | 54.96 | 31.11 | >72.00 | 49.38 |
| 180bp | 16,197,631 | **0.59** | **3.36** | **0.51** | 6.75 | 1.19 | 5.78 | 22.64 | 14.60 | >72.00 | >41.29 |
|  | 37,836,905 | **1.36** | **3.36** | **1.11** | 6.75 | 4.53 | 13.43 | 37.37 | 33.32 | >72.00 | >41.29 |
|  | 86,976,737 | **2.87** | **3.36** | **2.02** | 6.75 | 8.42 | 30.82 | 117.64 | 73.08 | >72.00 | >41.29 |

Next, we examine the scalability of different *k*-mer sets. We use 86,976,737 reads of 180bp to evaluate different batches of signatures, which are designed to test the worst scenario. Table 2 shows that TahcoRoll outperforms others in either time or memory in 13 out of 16 cases (indicated by red †). Compared the running time of “small” *k*-mers batches against other batches, TahcoRoll is more efficient in longer *k*-mers and *k*-mers with a wide range of *k*. This is due to less collision in those sets. Under a severe condition where there is a large number (12 and 24 million) of signatures with uniformly distributed characters, TahcoRoll requires more memory than MSBWT in five of these cases. However, it consistently requires half of the memory needed in Plain AC. It is worth mentioning that both MSBWT and KMC3 write a huge amount of intermediate files to disk (at least 16GB for MSBWT and 43GB for KMC3 in this dataset) to alleviate the memory bottleneck. In contrast, TahcoRoll is an in-memory approach that does not generate any intermediate data.

**Table 2:**
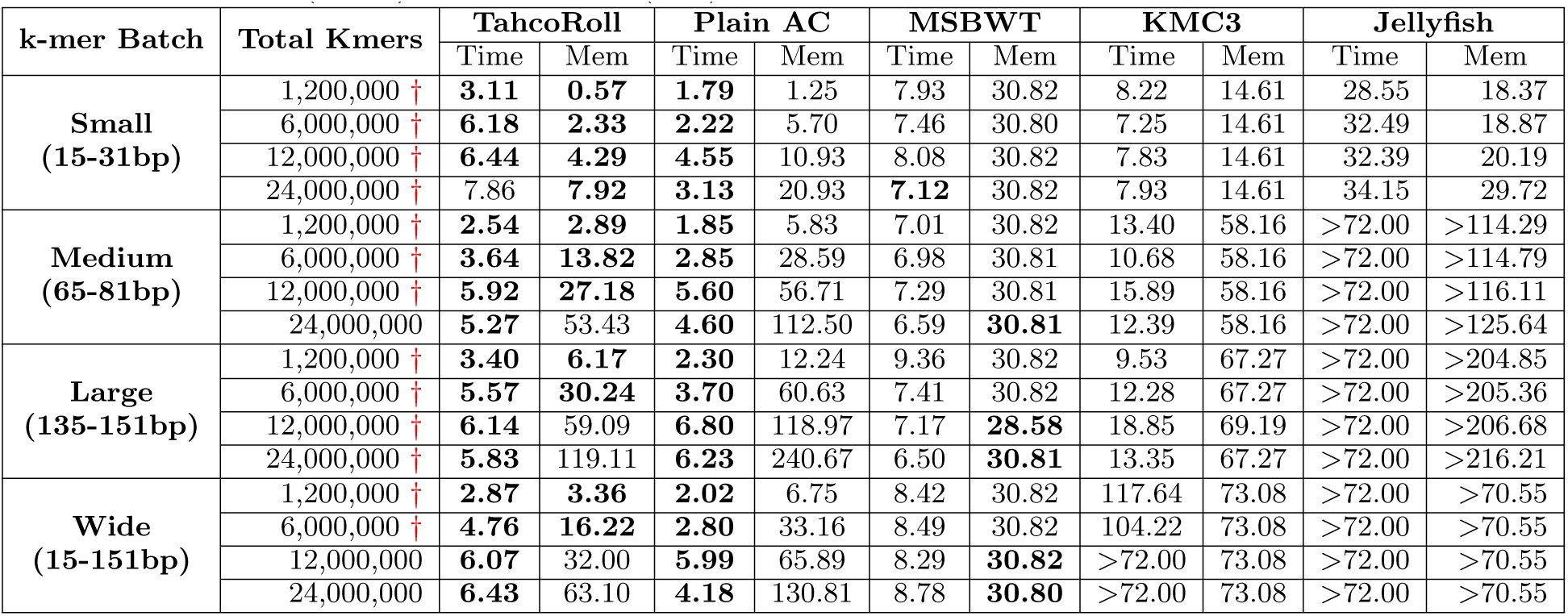
Time (Hour) and Memory (GB) of Synthetic Datasets over Different *k*-mer Sets

### 4.5 Real Datasets from Different Sequencing Platforms

The synthetic studies demonstrate the worst case of signature sets. Here, we examine the practical usage by analyzing signatures generated from real DNA sequences with six datasets from different sequencing platforms. The experiments are conducted on a desktop Linux machine of Core^*tm*^ i7- 3770 CPU@3.4.0GHz. Table 3 summarizes the nature and results of each dataset. From our previous studies, Jellyfish takes a long time (more than three days) to finish when we increase the read length, the number of reads, or the range of signature lengths. Li *et al.* [20] also observed that when there is not enough memory, Jellyfish writes intermediate results to disk and later merges them, which can be quite slow. Therefore, it is removed from this study. We highlight the most memory efficient approach in each experiment, and the top two time-efficient approaches since the running time are quite close between TahcoRoll and Plain AC. When the difference is large, we compute the fold change of each measurement to those reported by TahcoRoll. MSBWT fails to process long reads as the read length is not uniformly distributed. In the Illumina datasets, it runs slightly faster than TahcoRoll when the read length is 75bp, but requires twice as much memory. However, when the read length increases to 262bp, MSBWT is 1.8 times slower than TahcoRoll. KMC cannot index long reads from Oxford Nanopore as the data exceeds the buffer size automatically set by the program. When analyzing both Illumina and PacBio data, it is 7 to 9 times slower than TahcoRoll, and uses up all the memory available on the machine (32G) for SRR1293902 and GSM1254204. Evidently, the Aho-Corasick algorithm is the only approach that can profile both short and long reads with variable-length *k*-mers. TahcoRoll runs faster than Plain AC in the Illumina dataset because its automaton is lighter in memory and contains fewer collisions. Across all long read experiments, TahcoRoll takes slightly more time than plain AC, but requires half the memory.

**Table 3:** Running Time and Memory Footprint of Real Datasets

| Dataset |  | SRR1293902 | SRR1293901 | GSM1254204 | SRR5951587 | SRR5951588 | SRR5951600 |
| --- | --- | --- | --- | --- | --- | --- | --- |
| Source |  | RNA-Seq | RNA-Seq | RNA-Seq | WGS | WGS | WGS |
| Platform |  | Illumina HiSeq 2000 | Illumina MiSeq | PacBio | Nanopore | Nanopore | Nanopore |
| Number of Reads |  | 38,278,052 | 9,524,186 | 3,239,918 | 205,685 | 171,398 | 161,148 |
| Average Read Length |  | 75 | 262 | 1113 | 3kb | 8kb | 12kb |
| Time (Hour) | TahcoRoll | **1.57** | **1.64** | **1.47** | **0.31** | **0.55** | **0.81** |
|  | Plain AC | 1.74 | **1.71** | **1.40** | **0.28** | **0.45** | **0.69** |
|  | MSBWT | **1.47** | 3.01 (1.8X) | NA | NA | NA | NA |
|  | KMC3 | 12.46 (7.9X) | 12.31 (7.5X) | >13.82 (9.4X) | NA | NA | NA |
| Memory (GB) | TahcoRoll | **4.49** | **4.49** | **4.49** | **11.11** | **11.11** | **11.11** |
|  | Plain AC | 10.26 (2.3X) | 10.25 (2.3X) | 9.88 (2.2X) | 23.70 (2.1X) | 23.70 (2.1X) | 23.70 (2.1X) |
|  | MSBWT | 16.27 (3.6X) | 10.95 (2.4X) | NA | NA | NA | NA |
|  | KMC3 | 24.84 (3.6X) | >27.11 (>6.0X) | >26.65 (>5.9X) | NA | NA | NA |

## 5 Conclusions

In this paper, we propose a novel task of variable-length *k*-mer profiling. While the necessity of diversifying *k*-mer lengths has already been demonstrated in many studies [6,16,29], most of the existing works only support fixed-length *k*-mers and need an enormous amount of memory, disk space, and time to profile *k*-mers with a wide range of *k*’s. By leveraging the techniques of binarization and rolling hash for Aho-Corasick automaton, we construct a <u>t</u>hinned <u>Ah</u>o-<u>Co</u>rasick automaton accelerated by <u>roll</u>ing hash (TahcoRoll) to profile variable-length *k*-mers in genomic data. The main advantage of TahcoRoll is its in-memory property which does not require any disk space. A pilot study first gives a comprehensive overview of the comparisons of 14 approaches. Additional experimental results show that TahcoRoll scales well with both longer reads and a larger number of reads, especially that its memory usage is independent of the read data. It is the only approach that can efficiently process data from different sequencing platforms. In the evaluation of different *k*-mer sets, Aho-Corasick approaches use less time than others in most cases. Among the two implementations, TahcoRoll requires half of the memory needed in Plain AC.

Although all of our experiments focus on counting the frequency of a set of *k*-mers, the structure of this thinned automaton can be expanded to store essential information for each *k*-mer, such as its explicit positions in a genome for occurrence profiling. TahcoRoll can be further parallelized in two aspects. One is to separate the signatures automatically into a constant number of independent groups. These smaller set of signatures can be processed individually, leading to a thinner automaton. Although this requires TahcoRoll to scan the reads multiple times, the time complexity is still linear. The other aspect of parallelization is to perform read scanning with multiprocessing. Reads can be distributed into different threads. Multiple matching processes can access the thinned automaton in a shared memory.

TahcoRoll opens up the opportunity to profile a set of variable-length *k*-mers, especially in long read datasets. In the future, we plan to explore applying TahcoRoll in different sequencing applications, such as error correction for long read sequencing and discovering epigenetic marks through Bisulfite-Seq, to address various challenges in computational biology.

## Footnotes

1 https://pypi.python.org/pypi/pyahocorasick/

## Supplementary Materials

### A Proofs of Propositions in Section 2.3

Here we give the proofs of propositions shown in Section 2.3.

#### A.1 Proposition 1

*Proof*. Given the prefix length *i* and the number of possible characters *c*, there are *c^i^* possible prefixes in total. Under the assumption of uniform distribution, the probability that a particular prefix exists in *n* signatures is:

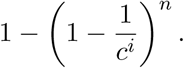

Therefore, the expected number of collided prefixes is:

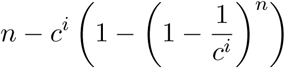

and the expected number of prefixes without any collision is:

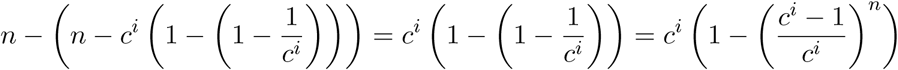

However, the expected number above includes the cases that fail before reaching the *i*-th character. Hence, the expected number of these cases should be deducted from the above number. Finally, the expected number of signatures that fail to find their length-*i* prefixes along the trie during its insertion is:

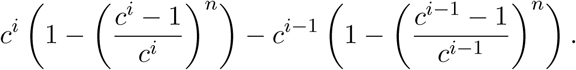

#### A.2 Proposition 2

*Proof*. Denote the expected number of signatures that fail to find their length-*i* prefixes on the trie during its insertion as *f*(*i*). Given the length of signatures *m*, each signature that fails to find the length-*i* prefix along the trie during its insertion will result in the addition of *m* − *i* + 1 nodes. From Proposition 2, the expected number of nodes in the trie is:

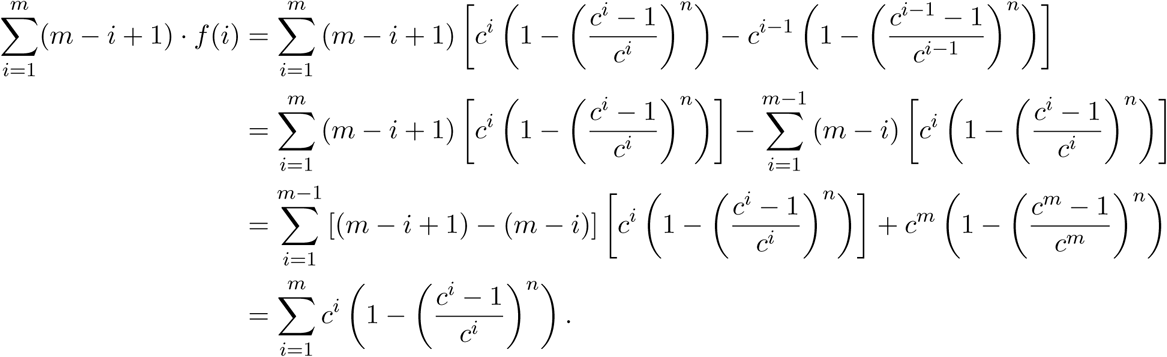

#### A.3 Proposition 3

*Proof*. Suppose that the number of signatures to be added into a trie is extremely large. The expected number of nodes with *c* possible characters is:

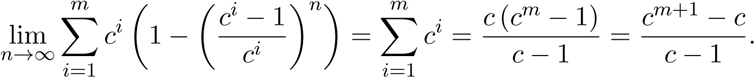

For the plain AC, there are four possible characters, i.e., `A`, `C`, `G` and `T`. Hence, the expected number of its nodes *N_A_* is:

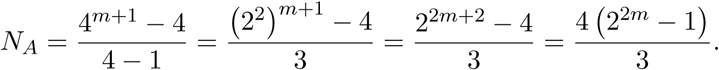

For the thinned automaton, there are two possible characters, i.e., `0` and `1`. Hence, the expected number of its nodes *N_T_* is:

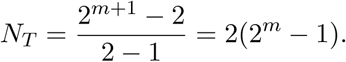

Finally, we compute the ratio of the expected number of two approaches as follows:

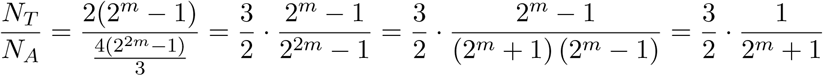

### B Detailed Formation of Rabin-Karp Rolling Hash

Formally, the hash value of a length-*L* input *t*[1… *L*] is defined as follows:

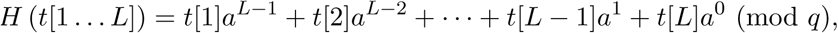

where *t*[*i*] is the *i*-th character of the input; *a* is a constant multiplier; *q* is a constant prime modulus. Based on the definition, the hash value of a length-*i* prefix of *t* can be recursively calculated with only the hash value of the length-(*i* – 1) prefix as follows:

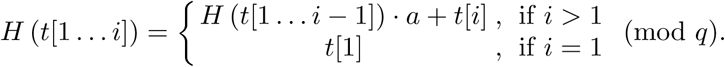

With bottom-up computation, hash values of all prefixes *H* (*t*[1… *i*]) can be preprocessed in both *O*(*L*) time and space complexity. Given the hash values of all prefixes, the hash value of any substring *t*[*i*… *j*] can be derived in *O*(1) as follows:

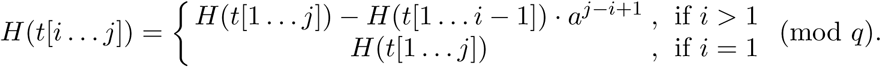

### C Related Work and Software Configurations

Existing *k*-mer counters index the read sequences into a compact and searchable structure, such as a hash table, a burst trie, or a compact suffix array. The occurrences of a specific *k*-mer can be retrieved by querying these data structures. Most of these counters are designed to process reads with fixed-size *k*-mers; several of them restrict the choice of *k* to fall within a threshold. These algorithms can be adapted to count *k*-mers of different sizes by repeating the process with different *k*’s. Here, we discuss different hashing and tree-based approaches, along with two baseline implementations.

#### C.1 Thread-Safe Shared Memory Hashing

**Jellyfish** is one of the early developed softwares to efficiently count the *k*-mer occurrences in read data. It exploits the CAS (compare-and-swap) assembly instruction to update a memory location in a multi-threaded environment, and uses the ‘quotienting technique’ and bit-packed data structure to reduce wasted memory. **Squeakr** builds an off-the-shelf data structure based on counting quotient filter (CQF). It maintains both global and local **CQF**s to facilitate updates of each thread. Both of these implementations provide an efficient function to query the counts of a specific *k*-mer.

Jellyfish provides the compilation of bindings to scripting languages, which allows us to query the output directly from Python. We first load the list of *k*-mers into memory. The indexing step is repeated for different *k*’s. One technique to retrieve *k*-mer frequency is to directly query the appropriate *k*-mer index through its querying function, denoted as *Jellyfish_query*. The other technique, denoted as *Jellyfish_dump*, is to dump the index into a human readable format, and scan through the result once. We implement both techniques for comparison.

#### C.2 Disk-Based Hashing

Disk-based hashing reduces the memory usage with complementary disk space. In general, this approach splits *k*-mers into bins, and stores them in files. Each bin is then loaded into the memory for counting. **DSK** divides *k*-mers into bins using a specific hash function based on the targeted memory and disk space. **KAnalyze** accumulates *k*-mers, and writes to disk when the allocated memory is full. **MSPKmerCounter** proposes a novel technique, Minimum Substring Partitioning, to reduce the memory usage of storing *k*-mers. One of the pitfalls is that **MSPKmerCounter** can only index reads with an odd number less than 64. **KMC**, **KMC2**, and **KMC3** are serial developments of a parallel *k*-mer counter. These methods scan reads one block at a time, and use a number of splitter threads to process these blocks. KMC2 leverages the concept of minimizer from MSPKmerCounter to further reduce disk usage. KMC3 accelerates the running time and optimizes the memory usage by taking a larger part of input data and better balancing the bin sizes.

##### DSK

To account for arbitrary *k*-mer lengths, we compile its source code with different parameters based on the range of the *k*-mer sizes. We set *cmake -DKSIZE_LIST* to be 32, 96, and 160 for small, medium, and large batches respectively. For the wide batch, we set this parameter to - *DKSIZE_LIST=”32 64 96 128 160”*. When running an experiment, we first load the *k*-mers into memory and determine the range of the sizes. We index the reads with different *k*’s using the main program of the appropriate compiled code, and dump the result into a human readable format with *dsk2ascii*.

##### KAnalyze

Similar to Jellyfish and DSK, we first load the list of *k*-mers into memory. KAnalyze provides two modes, *count* and *stream*. We use the *count* mode as it outputs a sorted tab-delimited file of **k*-mers* and their counts.

##### MSPKmerCounter

The software *MSPKC* contains three functions to be run in sequence: *partition, count,* and *dump.* The partition step divides short reads data using minimum substring partitioning, the count step computes the frequencies of existing *k*-mers, and the dump step converts the results into a human readable format. This sequel is repeated for each *k*.

##### KMC3

We use its main program to count the *k*-mer of all sizes seen in the list, with one *k* at a time. We set the -*cs* parameter to be 4294967295 to ensure all of the frequently occurred **k*-mers* are included. KMC provides an C++ API to load the compressed **k*-mer* occurrences into memory, and to retrieve the frequency of a specific *k*-mer. We develop a C++ module to utilize this API.

#### C.3 Probabilistic Hashing

To avoid counting *k*-mers with sequencing errors, **BFCounter** uses Bloom filter to identify all *k*-mers that are present more frequently than a threshold with a low false positive rate. The algorithm scans read data in two passes to avoid reporting false positive counts, and is designed to handle *k* less than 64. **khmer** uses a streaming-based probabilistic data structure, CountMin Sketch [9]. The algorithm is designed to perform in-memory nucleotide sequence *k*-mer counting, and cannot handle *k* larger than 32.

##### BFCounter

Similar to DSK, BFCounter provides different configurations to comprehend larger **k*-mers.* We compile three versions based on the largest *k* in the dataset. We set *MAX_KMER_SIZE* to 32 for the small batch, 96 for the medium batch, and 160 for both of the large and wide batches. The count function of BFCounter requires an estimation of the number of **k*-mers*, and we use *KmerStream* [23] to pre-compute the **k*-mer* statistics in read data. We use the *dump* function to convert the results into a human readable format to extract the frequencies.

##### khmer

We use its Python wrapper script, *load-into-counting,* to perform counting, which writes a *k*-mer graph for each *k* to files. We repeat this step for different *k*’s. Each *k*-mer graph is loaded back to the memory one at a time, allowing us to query the count. We set the maximum amount of memory for the data structure to be 10G as the required parameter. However, increasing this parameter does not alleviate the limitation of using larger *k*.

#### C.4 Suffix-Arrays

Suffix-arrays are powerful data structures that present the potential of searching arbitrary *k*-mers without any restriction of *k* on a single scan. However, constructing a suffix-array on read data can be computationally expensive. **Tallymer** is tailored to detect *de novo* repetitive elements with sizes ranging from 10 to 500bp in the genome. The algorithm first constructs an enhanced suffix-array, and indexes *k*-mers for a fixed value of *k*. The indexing step needs to be repeated for different *k*’s. **MSBWT** is designed to compress raw reads via a multi-string variant of Burrows-Wheeler Transform (BWT). Instead of concatenating all reads and sorting, MSBWT builds a BWT on each string and merges these multi-string BWTs through a small interleave array. The final structure allows a fast query of *k*-mers of arbitrary *k*.

##### Tallymer

Tallymer builds an enhanced suffix-array on read data through the function *gt suffixerator*. Since it provides a function to query the count of a set of *k*-mers, we first separate our representative *k*-mers into different files based on their sizes *k*. For each *k*, we use *gt tallymer mindex* to extract the *k*-mer index and count from the enhanced suffix-array, and use *gt tallymer search* to retrieve their counts.

##### MSBWT

We build the multi-string BWTs on all reads. This compressed representation is stored in files, and later loaded in memory for frequency retrieval. According to the complexity analysis provided by [15], the construction time is *O*(*l*× *LCS*) where LCS is the longest common substring between any two BWTs, and the search is linear to each *k*-mer size, which is O(M) for all *k*-mers.

#### C.5 Burst Tries

A common drawback of hash-based and tree-based solutions is that they can suffer from huge cache misses when handling large datasets. **KCMBT** uses a cache efficient burst trie to store compact *k*-mers. The trie structure stores *k*-mers that share the same prefix in the same container. When a container is full, *k*-mers are sorted and burst. A good balance between the container size and the tree depth is essential to avoid constant sorting and bursting. As a result, KCMBT uses hundreds of trees. Unfortunately, it is limited to process *k*-mers with *k* less than 32.

##### KCMBT

We first load the list of *k*-mers into memory. KCMBT generates binary files containing *k*-mers and their counts. We use *kcmbt_dump* to convert the binary data into human readable files. We scan through each *k*-mer in the output files to extract the frequencies of our *k*-mers.

#### C.6 Naïve Implementation

The naïve implementation is to scan through the reads multiple times with different window sizes. A list of representative *k*-mers is loaded into memory using a hash table, where the key represents a *k*-mer and the value stores its occurrence. Each read is scanned *d* times where *d* is the number of different sizes we observed in the list. Assume that the total length of read sequences is *l*, and the total length of *k*-mers is *M* (*l* >> *M*), the time complexity for constructing the hash is *O*(*M*) and for scanning reads and querying the hash table is*O*(*l* × *k*) for one window size *k*. Considering the *k*-mer size ranges from *k*_1_ to *k*_d_, then the total examination time is 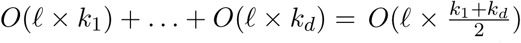. Theoretically, the naïve implementation is light in memory since we only store the *k*-mers, but requires an extensive running time.

#### C.7 Plain Aho-Corasick

We use one of the finest and fastest Aho-Corasick libraries in Python, *pyahocorasick* ^1^, as our benchmark to access the speed and memory usage for this approach. One drawback of this library is that the automaton trie needs to be constructed and finalized ahead of time before the search, and thus the frequency of each *k*-mer cannot be stored in the trie. Consequently, we use an additional structure to store the *k*-mers along with their frequencies in read data. Although this approach scan through reads in linear time, the memory footprint is expected to be large as we increase the number and size of our *k*-mers.

